# Abiotic and biotic ecological factors affect the genomic consequences of plant-butterfly coevolution

**DOI:** 10.64898/2026.09.23.753697

**Authors:** Tyler Figueira, Quint Rusman, Florian P. Schiestl

## Abstract

Variation in abiotic and biotic factors drives unique coevolutionary trajectories, yet we know little about how individual and interactive effects from such environmental factors alter the genomic basis of coadaptation. As a follow-up study after 6 generations of experimental coevolution, we analyzed whole-genome re-sequencing data from “fast-cycling” *Brassica rapa* plants that coevolved with a pollinating herbivore (*Pieris rapae* butterflies) in two different temperature environments (ambient/elevated) and in the presence/absence of bumblebees that did not coevolve. We found that genomic divergence between plant populations coevolving in the presence or absence of bumblebees was larger relative to plant populations coevolving in different temperature environments. Next, we found that the presence of bumblebees affected the number of genomic regions under selection differently depending on which temperature environment plants coevolved in. More specifically, we found no genomic regions under selection when plants coevolved in elevated temperatures and in the absence of bumblebees, but the most genomic regions under selection when plants coevolved in elevated temperatures and in the presence of bumblebees. Finally, genomic evolution underlying defense traits was larger when plants coevolved in the presence of bumblebees relative to when plants coevolved in the absence of bumblebees, which fits the prediction of coevolution being more antagonistic in the presence of a non-herbivorous co-pollinator. Overall, our results highlight how modifications in temperature and community context, potentially stemming from global environmental change, will affect the genomic basis of coevolution.

## 1 Introduction

Coevolution is reciprocal evolutionary change between interacting species (Thompson, 2005), driving the evolution of biodiversity at both the phenotypic and genetic levels (Bento et al., 2017; Edger et al., 2015; Eizaguirre & Lenz, 2010; Hoso et al., 2010). When coevolution differs between populations of interacting species, coevolution can promote divergence (Friman & Buckling, 2013; Laine, 2009; Parchman et al., 2016). Differences in abiotic and biotic factors across populations can drive such variation in coevolutionary interactions (Benkman, 1999; Thompson & Cunningham, 2002; Toju et al., 2011). For example, environmental factors can induce phenotypic plasticity in interacting species, potentially altering the fitness outcome of the interactions and/or the presence/strength of reciprocal selection (Rusman, Figueira, et al., 2025; Rusman, Traine, et al., 2025; Thompson, 2005). Whereas multiple studies have shown that environmental factors can alter phenotypic coevolution between interacting species (Benkman, 1999; Thompson & Cunningham, 2002; Toju et al., 2011; Toju & Sota, 2006), we know much less about how abiotic and biotic factors affect genomic coevolution. Such knowledge is necessary to understand under what conditions coevolution can promote divergence between populations.

Abiotic factors, including climatic factors such as temperature, have been shown to affect coevolutionary interactions in both experimental and field studies. For example, experimentally manipulated elevated temperatures reduced reciprocal selection between *Brassica rapa* plants and *Pieris rapae* butterflies, potentially because temperature-induced phenotypic plasticity disrupted trait-fitness relationships (Rusman, Traine, et al., 2025). Experimental bacteria-phage coevolution was weaker in constant elevated temperatures relative to both constant standard temperatures and fluctuating elevated temperatures because lower phage densities in constant elevated temperatures reduced encounter rates between bacteria and phages (Duncan et al., 2017). Conversely, warmer, more productive environments promoted coevolution between Japanese camellia trees (*Camellia japonica*) and their obligate seed weevil antagonists (*Curculio camelliae*) because trees could invest more resources to produce thicker seeds, thereby selecting for longer snouts in weevils (Toju et al., 2011; Toju & Sota, 2006). Overall, due to abiotic factors having large effects on phenotypic coevolution, it is expected that such factors will also have a considerable impact on genomic coevolution (Thompson, 2005).

Similar to abiotic factors, the presence of additional biotic interactions and variation in the community context can affect phenotypic coevolutionary trajectories. For example, the presence of red squirrels (*Tamiasciurus hudsonicus*) prevented coevolution between lodgepole pines (*Pinus contorta ssp. Latifolia*) and red crossbills (*Loxia curvirostra* complex) because squirrels generally harvest pinecones earlier in the year relative to crossbills (Benkman, 1999). The outcome of the interaction between *Lithophragma parviflorum* plants and a pollinating seed-predator, *Greya politella* moths, was more antagonistic when additional pollinators were present, driving increased defenses against seed herbivory from *Greya* moths (Thompson & Cunningham, 2002). While *Taricha granulosa* newts and *Thamnophis sirtalis* snakes generally show matched levels of toxin defense and resistance across populations (Brodie et al., 2002), newts that overlap geographically with more snake species and snakes that overlap geographically with more toxic newt species exhibit higher levels of defense and resistance, respectively, suggesting that the community context influences coevolution (Gilbert et al., 2023). Importantly, the assemblage of newt and snake species varied due to climatic variables, indicating that the co-occurrence of both abiotic and biotic variation combined to affect coevolution. This is particularly relevant to the current study of coevolution because global environmental change will influence both abiotic and biotic factors; associated elevated temperatures and a higher prevalence of extreme weather conditions will affect the distribution of interacting species (IPCC, 2018, 2023; Parmesan, 2006; Parmesan & Yohe, 2003; Rubenstein et al., 2023). Therefore, to understand how global environmental change will impact coevolutionary interactions, it is important to study both abiotic and biotic factors together.

A powerful approach for examining how individual and interactive effects from multiple factors affect coevolution is experimental coevolution (Brockhurst & Koskella, 2013; Kawecki et al., 2012). Importantly, experimental coevolutionary studies can manipulate the potential for reciprocal selection/adaptation (Brockhurst & Koskella, 2013; Paterson et al., 2010; Schulte et al., 2010), thus providing stronger evidence for coevolution relative to field and phylogenetic comparative studies that infer coevolution from trait correlations (however see: Brodie & Ridenhour, 2003). Experimental coevolution studies on bacteria and phages showed that the community context (monoculture vs polyculture) affected the evolution of phage resistance in bacteria (Castledine et al., 2024), while elevated temperatures promoted host persistence and parasite extinction, thereby altering coevolution (Zhang & Buckling, 2011). A recent coevolution experiment investigated how elevated temperatures and the presence of bumblebees affected coevolution between *Brassica rapa* plants and *Pieris rapae* butterflies (Figueira et al., *in review*; Rusman, Figueira, et al., 2025; Rusman, Traine, et al., 2025). Both single-factor effects of elevated temperatures and the addition of bumblebees resulted in increased antagonistic coevolution (i.e., increased evolution of herbivore resistance/deterrence in plants), whereas the combination of elevated temperatures and bumblebees led to reduced antagonistic coevolution (Rusman, Figueira, et al., 2025). Overall, experimental coevolution is a promising approach to isolate how environmental factors alter coevolution, yet it has not been used to examine how abiotic and biotic factors interact to affect the genomic consequences of coevolution.

Here, we analyzed the genomes of plants from this recent coevolution experiment that investigated how coevolution was affected by different temperatures and pollinator-community contexts (Figueira et al., *in review*; Rusman, Figueira, et al., 2025; Rusman, Traine, et al., 2025). In this experiment, “fast-cycling” *Brassica rapa* plants coevolved in response to 6 generations of selection where they interacted with coevolving *Pieris rapae* butterflies in either “ambient” (23±2°C) or “elevated” (27±3°C with 24h of 30±3°C one day per week) temperature environments and in the presence/absence of non-coevolving bumblebees (*Bombus terrestris*). In natural conditions, *P. rapae* is a generalist flower visitor that pollinates *B. rapa* and oviposits on plants that the Brassicaceae-specialized caterpillars feed on (Feltwell, 1981; Traine et al., 2024; Wheat et al., 2007). Previous experimental work assessed the phenotypic consequences of the plant-butterfly interaction by measuring traits related to both mutualistic (pollinator attraction/butterfly foraging) and antagonistic (defense/counterdefense) components of coevolution (Figueira et al., *in review*; Rusman, Figueira, et al., 2025; Rusman, Traine, et al., 2025), ultimately providing strong evidence of coevolution between plants and butterflies. In this study, we focused on the plant side of the coevolutionary interaction and tested whether abiotic and biotic factors: 1) differ in their effect on genomic divergence, 2) interact to affect patterns of genomic adaptation, and 3) interact to affect the strength of genomic evolution underlying attraction and defense traits. We predicted that: 1) the effect of bumblebee presence on genomic divergence will be larger than that of elevated temperatures because previous studies have shown biotic pollination to be a strong driver of genomic divergence (Busch et al., 2022; Figueira et al., 2025; Frachon et al., 2023), 2) the effect of bumblebee presence on genomic adaptation will depend on the temperature environment - more specifically, plants that coevolved with butterflies in stressful elevated temperatures with bumblebees will have less genomic regions under selection relative to plants that coevolved in ambient temperatures with bumblebees, and 3) the strength of genomic evolution underlying traits associated with defensive functions will be higher for plants that coevolved with butterflies in the presence of bumblebees because coevolution with a pollinating herbivore can be more antagonistic for the plant with the addition of mutualistic bumblebee pollinators (Rusman, Figueira, et al., 2025; Thompson & Cunningham, 2002).

## 2 Materials and Methods

### 2.1 Origin of plant samples

The phenotypic and genomic data used in the current study are derived from the final generation of *Brassica rapa* plants that evolved over the course of a coevolution experiment (Rusman, Figueira, et al., 2025); the methodology from this previous experiment is summarized in section 2.2. In short, plants evolved over 6 generations in response to different combinations of two temperature environments and three biotic treatments (interactions with coevolving butterflies, interactions with coevolving butterflies and non-coevolving bumblebees, and no biotic interactions - plants were randomly hand-pollinated as a control). After 6 generations of selection, plants from all treatments were grown without any insects to minimize maternal effects (generation 7), and plants from generation 7 were hand-pollinated. The seeds from these crosses were the 8^th^ and final generation. 8^th^-generation plants were grown to assess phenotypic and genomic evolution (Rusman, Figueira, et al., 2025), and the data used in this study was collected from these 8^th^-generation plants. The methodology used in the current study begins at section 2.3.

### 2.2 Experimental (co)evolution summary

“Fast cycling” *B. rapa* plants evolved in response to two temperature environments and three biotic treatments for 6 generations in greenhouses with controlled conditions (Rusman, Figueira, et al., 2025). Plants evolved in either ambient temperatures (23±2⁰C) or elevated temperatures (27±3⁰C with 24h of 30±3⁰C once per week). The ambient and elevated temperature environments mirrored current and future mean temperatures of late spring/early summer in northern Switzerland predicted by the CH2018 model (Fischer et al., 2022). For biotic treatments, plants either 1) coevolved with coevolving butterflies (*Pieris rapae*) in the absence of bumblebees (*Bombus terrestris*), 2) coevolved with coevolving butterflies in the presence of non-coevolving bumblebees, or 3) evolved without biotic interactions and were hand-pollinated (control treatment). Coevolving butterflies were kept in greenhouse cabins with their respective temperature environment (ambient or elevated); see Rusman et al. 2025 for detailed information on the rearing of the butterfly evolution lines. There was no gene flow between replicates or treatments, allowing replicates to represent independent population units. Importantly, within each treatment-replicate combination, plants and butterflies were allowed to adapt to each other: each plant and female butterfly contributed offspring to the next generation proportionally to its total number of offspring (i.e., fitness), thus allowing reciprocal adaptation (Figueira et al., *in review*; Rusman, Figueira, et al., 2025; Rusman, Traine, et al., 2025).

#### 2.2.1 Plant and butterfly starting populations

We used “fast cycling” *B. rapa,* bred for short generation times and ample genetic variation (Wisconsin Fast Plants standard variety seeds, Carolina Biological Supply, United States of America). “Fast cycling” *B. rapa* plants are annual and mostly self-incompatible. In natural and agricultural conditions, conspecific *B. rapa* plants interact with both mutualistic pollinators and antagonistic herbivores, making them an attractive study system (Atmowidi et al., 2007; Lamb, 2026; Rader et al., 2009).

We established the starting plant population in 2020. We first created full-sib plant families by growing 300 *B. rapa* plants and assigned them to pairs. Within each pair, five flowers from each plant were reciprocally hand-pollinated. The resulting seeds from both parents were mixed to form one full-sib family. Thus, 150 full-sib plant families were created, and we used 98 of them, with 49 unique families randomly assigned to two replicates.

To grow plants for the experimental coevolution study, seeds were sown into multi-pot trays filled with a mixture consisting of 4 equal parts: compost (3 years old; Zurich Botanical Garden, Switzerland), peat (medium structure; Zelta Zeme, Latvia), pumice (1-5 mm; Ökohum GmbH, Switzerland), and sterilized sieved topsoil (Zurich Botanical Garden, Switzerland). Seed trays were placed in phytotrons (22±1⁰C, 60%RH, 24h light) for 8 days. Germinated seedlings were repotted into individual pots (7×7×8 cm), filled with standardized soil (Patzer-Erden GmbH, Sinntal, Germany), and placed into greenhouses with their respective temperature environment (ambient: 23±2⁰C or elevated: 27±3⁰C with 24h of 30±3⁰C one day per week) and similar other greenhouse conditions (60%RH, 16h:8h, light: dark).

To establish a butterfly starting population, wild *P. rapae* butterflies and eggs were collected from 6 locations. This includes 5 sites in Switzerland (Binz, Dübendorf, Einsiedeln, Sonnental, Zürich) and 1 site in the Netherlands (Renkum). Each Swiss population was crossed with the Dutch population by placing virgin females of each individual Swiss population together with virgin males of the Dutch population, and vice versa. The hybrid populations from each Swiss and Dutch population crossing were combined (Dutch female X Swiss male + Swiss female X Dutch male) to create 5 hybrid lines (BinzXNL, DübXNL, EinXNL, SonXNL, ZürXNL).

Finally, 3 rounds of crossings between hybrid lines were performed to create a single population with genetic material from the 6 sampled populations. Adult butterflies from the starting population were kept in a rearing cage (90 x 60 x 60 cm) with water, nectar, and flowering “fast cycling” *B. rapa* plants to oviposit on. Emerging young larvae (L1-L3) fed on *B. rapa* plants, while late-instar larvae (L4-L5) fed on savoy cabbage leaves (*B. oleraceae* var. sabauda).

#### 2.2.2 Biotic pollination and herbivory

Each generation, butterflies and bumblebees were used in pollination bioassays. Butterflies and bumblebees had no contact with experimental plants before bioassays. Newly eclosed butterflies mated 2 days before bioassays and were starved 1 day before bioassays. Two bumblebee (*B. terrestris*) hives were purchased from Andermatt Biocontrol for each generation (Hummelvolk Bombus Maxi, Andermatt Biocontrol, Grossdietwil, Switzerland). Bumblebees fed on flowering “fast cycling” *B. rapa* plants with supplementary nectar (BioGluc, Biobest, Westerlo, Belgium) and pollen (Blütenpollen Multiflora, KoRo Handels GmbH, Berlin, Germany). To avoid pollen from standard *B. rapa* plants mixing with plants from evolutionary treatments (pollen contamination), plants were removed from bumblebee cages 2 days before bioassays. Bumblebees were starved 16 hours before bioassays.

Pollination bioassays took place 24-27 days after sowing, where butterflies interacted with plants in the presence or absence of bumblebees and inside greenhouses with the correct temperature environment. Plants were placed into a seven-by-seven matrix inside a flight cage (2.5×1.8×1.2m). For plants that coevolved in the absence of bumblebees, 25 mated female butterflies were released into their respective flight cages and recaptured after 2 full days. For plants that coevolved in the presence of bumblebees, 20 mated female butterflies were released into flight cages for 2 days, and simultaneously, 6 worker bumblebees were released into flight cages sequentially. Each bee was recaptured after it visited 4 plants.

1-3 days after butterflies were recaptured, the number of eggs oviposited on each plant was counted. Plants were ranked based on the number of eggs that they received. For example, plants ranked at 1 received the most eggs in the treatment-replicate combination, while plants ranked at 49 received the least number of eggs in the treatment-replicate combination. Based on rankings and 7-8 days after oviposition, plants were assigned one of five herbivory treatments (0-4 caterpillars left on plants) by removing unhatched eggs and excess larvae. More specifically, 4 caterpillars were left on plants ranked from 1 to 9; 3 caterpillars were left on plants ranked 10 to 19; 2 caterpillars were left on plants ranked 20 to 29; 1 caterpillar was left on plants ranked 30 to 39; and all caterpillars were removed for plants ranked 40 to 49. The range of 0 to 4 caterpillars per plant simulated natural levels of herbivory while reducing plant mortality. For example, minimum values of larval survival may reach 0.1% (Parker, 1970), while plants struggle to survive due to herbivory from more than 4 caterpillars (Rusman, Traine, et al., 2025; Traine et al., 2025).

10-11 days after oviposition, checks were performed to ensure that all plants had the correct number of caterpillars. At the same time, plastic cones were placed around plants to prevent late-instar larvae (L4-L5) from migrating between plants. Plastic cones were made with 3 PVC A4-sized pages and lined the top border with cotton wool.

3-5 weeks after oviposition, pupae were collected and stored in refrigerators (10⁰C). At the same time, plants were moved to a different greenhouse without watering. Once the fruits dried sufficiently, the number of fruits per plant was counted. The number of seeds per plant was counted by extracting the seeds from the fruits, placing all the seeds from a plant onto a white plate, photographing the plate, and using ImageJ. The contribution of each plant to the next generation was calculated with this formula: 49/(population sum of seeds/individual number of seeds).

#### 2.2.3 Control treatment pollination

24-27 days after sowing, plants from the control treatment were also pollinated. Within each replicate, pollen was collected from 3-5 flowers of each plant with a makeup brush. 6 flowers per plant were pollinated with the pollen mix. All successfully reproducing plants always contributed one individual seed to the next generation, while plants that did not successfully reproduce (i.e., due to mortality, fertility, or germination-related mechanisms) were replaced randomly by a successfully reproducing plant.

#### 2.2.4 Generations 7 and 8

After 6 generations of selection, plants from all treatments were grown without any insects to minimize maternal effects (generation 7). Plants from generation 7 were hand-pollinated following the protocol as described above, except that 10 flowers were pollinated instead of 6. The seeds from these crosses were the 8^th^ and final generation. These 8^th^-generation plants were then grown for an experiment to assess phenotypic and genomic evolution.

#### 2.2.5 Experiment to assess evolution summary

36 plants from randomly chosen seed families of the 8^th^ and final generation of each treatment-replicate combination were grown and phenotyped as described in Rusman, Figueira, et al., 2025. Greenhouse conditions and sowing and potting followed the same protocols as described earlier. However, plants and butterflies were only grown in the ambient temperature environment (ambient: 23±2⁰C) to prevent plasticity due to elevated temperatures. Treatments were assigned to one of four cohorts separated by two weeks to synchronize the developmental time of plants and butterflies. For bioassays, 18 newly eclosed female butterflies were collected from cages of each treatment-replicate combination.

#### 2.2.6 Phenotyping

Between 21-25 days after sowing, plants were phenotyped (see here for more detailed methodology: Figueira et al., *in review*; Rusman et al., 2025; Traine et al., 2026). In short, morphological traits such as plant height, the number of flowers and leaves, and flower size were measured. The amount of nectar per plant and the emission of 14 floral volatile compounds were quantified. ∼15 mg of fresh leaf tissue was sampled to measure the concentration of 14 glucosinolate secondary metabolites. Leaf tissue was taken before herbivory and then wrapped in foil and placed into liquid nitrogen.

#### 2.2.7 Herbivory treatment

29-32 days after sowing, butterflies were allowed to interact with plants from their respective treatment-replicate combinations. Flowering plants (range of open flowers: 0-37) were randomly arranged in a six-by-six matrix inside a flight cage (2.5×1.8×1.2m). 18 female butterflies per treatment-replicate combination were marked and released into the flight cage containing plants from their respective treatment-replicate combination. Butterflies could interact with plants for 2 days between 10:00-12:00 and 13:00-16:00 (10 hours total, 5 hours per day).

1-3 days after the second day of bioassays, the number of eggs oviposited on each plant was summed. Plants were ranked depending on the number of eggs that they received: rank 1 indicated that the plant received the maximum number of eggs within the treatment-replicate combination, and rank 36 indicated that the plant received the minimum number of eggs within the treatment-replicate combination. Based on the plant’s rank, they were assigned to one of five herbivory treatments. 7-8 days after oviposition, unhatched eggs and excess larvae were removed until plants had the appropriate number of caterpillars (range of 0-4 caterpillars left on the plant). More specifically, 4 caterpillars were left on plants ranked 1-7, 3 caterpillars were left on plants ranked 8-14, 2 caterpillars were left on plants ranked 15-21, 1 caterpillar was left on plants ranked 22-28, and all caterpillars were removed from plants ranked 29-36.

7-8 days after oviposition, we also recorded whether plants induced the hypersensitive response (HR). A plant was considered to have induced HR when a ring of black necrotic tissue surrounding the base of an egg was observed (Fatouros et al. 2014). Per plant, the proportion of HR induction was calculated as the number of eggs with black necrotic rings surrounding the base, divided by the total number of oviposited eggs.

### 2.3 Methodology used in the current study

#### 2.3.1 DNA extractions

During the experiment to assess evolution (21-25 days after sowing), ∼15 mg of fresh leaf tissue was sampled from plants of the 8^th^ generation for genomic analyses. As mentioned earlier, plants were only grown in the ambient temperature environment, and leaf tissue was taken before herbivory. Leaf tissue was wrapped in foil and placed into liquid nitrogen.

Additionally, after the experiment, leaf tissue was sampled in the same way from plants of generation one (G1). In total, leaf tissue was sampled from 648 plants for genomic analyses (individuals per treatment and replicate = min: 36, max: 72).

Before DNA extractions, we freeze-dried leaf tissue samples at −70°C and then ground samples into a fine powder with glass beads. We extracted DNA with a customized SBEADEX Mini Plant DNA Purification kit (Biosearch Technologies). We performed DNA extractions with a KingFisher Flex machine (ThermoFisher Scientific) at the Genomic Diversity Center of ETH (Zurich, Switzerland).

#### 2.3.2 Genotype assembly and filters

We shipped purified DNA to the Poland branch of the Beijing Genomics Institute (BGI; Shenzhen, China). BGI prepared libraries and performed resequencing and initial quality controls. We received ∼10Tb of clean, raw reads and then applied FastQC/multiqc to assess the quality of the fasta files. We used cutadapt to trim reads (Martin, 2011), aligned fasta files with the Burrows-Wheeler algorithm (BWA; v0.7.17; (Li & Durbin, 2009)), and used the *mpileup* function in bcftools for variant calling (Danecek et al., 2021). We removed indels and kept only biallelic SNPs.

#### 2.3.3 Alignment and SNP calling

We aligned fasta files against the Chiifu (v3.0) reference genome (genome size = ∼455 Mb, BRAD, http://brassicadb.cn; (Zhang et al., 2018)). We used VCFtools to filter samples with -- max-missing 90 and sites with --minGQ 15 and MAF – 0.01 (Danecek et al., 2011). The mean depth per site was ∼15, so we filtered for --min-meanDP 5 --max-meanDP 45. After filtering, there were ∼4 million SNPs, and the mean ± standard deviation coverage per sample was 16 ± 1.17 (min = 9.35, max = 20.04).

### 2.4 Data analyses

Unless otherwise noted, all analyses were run in R version 4.4.2 (R Core Team, 2022).

#### 2.4.1 *F_ST_* scans

To compare the strength of genetic differentiation between plants that evolved in different temperature environments and between plants that evolved in the presence/absence of bumblebees, we ran *F_ST_* scans with non-overlapping windows of 10,000 bp in VCFtools (Danecek et al., 2011). We ran *F_ST_* scans within replicates. To assess the effect of temperature, we grouped plants that evolved in the presence and absence of bumblebees in the ambient temperature environment and grouped plants that evolved in the presence and absence of bumblebees in the elevated temperature environment. To assess the effect of the presence/absence of bumblebees, we grouped plants that evolved in the presence of bumblebees in both temperature environments and grouped plants that evolved in the absence of bumblebees in both temperature environments. In both comparisons, each group comprised 72 plants (36 plants per treatment with 2 treatments).

To assess whether *F_ST_* values were higher due to evolving in different temperature environments, or due to evolving in the presence or absence of bumblebees, we ran a generalized linear mixed model with the glmmTMB package (Brooks et al., 2017). The response variable was the *F_ST_* value for a given 10kb window (n = 110667). The fixed effect was the comparison (temperature or bumblebee). We included replicate, chromosome, and starting base position of the 10kb window nested within the chromosome as random effects. The range of *F_ST_* values was: [0-1), thus, we used an ordinal beta family (Kubinec, 2023).

#### 2.4.2 Selection scans

We ran Cochran–Mantel–Haenszel (CMH) tests (Spitzer et al., 2020) and CLEAR: Composition of Likelihoods for Evolve and Resequence Experiments (Iranmehr et al., 2017) to identify SNPs with significant allele frequency change across both replicates. As CMH tests and CLEAR incorporate allele frequencies across all replicates (2 in this study), they have strong power to detect treatment-level effects. However, there is a trade-off as it prohibits the identification of replicate-specific evolution. CMH tests and CLEAR have high power and low false positive rates relative to other replicate-incorporating methods (Vlachos et al., 2019).

We ran CMH tests with allele frequencies of plants from generation one (G1) and plants from the final generation (G8). We used the R package ACER (Spitzer et al., 2020). To correct for multiple testing, we used the *p.adjust* function in R with the False Discovery Rate (FDR) method. We identified genomic regions under selection with the local score approach (Fariello et al., 2017), which sums selection signals from individual SNPs (*p*-values generated from CMH tests). We also ran CLEAR on data files in a sync format. For each treatment-replicate combination, we used the minimum number of plants that contributed to the next generation, across all generations, as a proxy for effective population size.

We considered a genomic region to be under selection when all 4 criteria were met: 1) the region was identified using the local score approach, 2) the FDR-adjusted *p*-value < 0.05 for the SNP with the lowest *p*-value generated from CMH tests inside the genomic region, 3) the selection coefficient generated from CLEAR was in the top 5% for the SNP with the lowest *p*-value generated from CMH tests inside the genomic region, and 4) the genomic region did not overlap with genomic regions under selection in the greenhouse control treatment for the respective temperature environment. If a genomic region met all 4 criteria, then we used the SNP with the lowest *p*-value from CMH tests for downstream genotype-phenotype associations.

Finally, to assess how temperature and bumblebee presence interacted to affect the number of selected genomic regions, we performed pairwise proportion tests between plants from all treatments except for plants from the control treatment. We compared the proportion of treatment-specific selected genomic regions over the total number of selected genomic regions across all treatments. We performed proportion tests with the *prop.test* function in R. We then corrected for multiple testing with the *p.adjust* function using the FDR method. We considered pairwise comparisons to be significant when the FDR-adjusted *p*-value < 0.05.

#### 2.4.3 Genotype-phenotype associations

To identify genotype-phenotype associations, we ran generalized linear mixed models with the *glmmTMB* or *betareg* package for each genomic region with signatures of selection (Brooks et al., 2017; Cribari-Neto & Zeileis, 2010). We ran genotype-phenotype associations for plants that evolved in different treatments separately and included control plants from their respective temperature environments in each of the runs. The average sample sizes for each run ± standard deviation are listed for plants from the different treatments: plants that evolved in ambient temperatures without bumblebees: 154.75 ± 6.43; plants that evolved in ambient temperatures with bumblebees: 125.11 ± 13.56; plants that evolved in elevated temperatures with bumblebees: 125.02 ± 17.89. For plants that evolved in ambient temperatures without bumblebees, we published data in an earlier study with genotype-phenotype associations for all traits except for glucosinolates (Figueira et al., *in review*). Thus, for the current study, we only ran genotype-phenotype associations for glucosinolate traits for plants from this treatment, but we display all recorded associations.

We used genotypes from SNPs with the lowest *p*-value from CMH tests within selected genomic regions to test for genotype-phenotype associations. We used bcftools *query* to extract genotypes from VCF files (Danecek et al., 2021), which returned genotypes for each plant in this format: (allele 1/allele 2). As we filtered the VCF files for biallelic sites, allele 1 and allele 2 were coded either as 0 or 1. A value of 0 indicates that the allele was the reference allele (the allele from the reference genome), and a value of 1 indicates that the allele was the alternative allele. Thus, all possible genotypes were either (0/0), (0/1), (1/0), or (1/1), and because we did not perform phasing, heterozygotes, coded as (0/1) or (1/0), were considered equal.

We then tested for genotype-phenotype associations across 37 traits (5 morphological traits, nectar amount, 14 floral volatile compounds, total floral volatile emission, 14 glucosinolate traits, total glucosinolate emission, induction of the hypersensitive response (HR; proportion of eggs that had a black necrotic ring around them over the total number of eggs that a plant received), and fruit damage. In all models, the phenotypic trait was the response variable, and the plant genotype was the fixed effect. However, we used different error distributions between traits. For the number of flowers, we used a negative binomial distribution. We used a Tweedie distribution for nectar amount and a Gaussian distribution for plant height, number of leaves, and flower size. We logged the values of floral volatile emissions and glucosinolate concentrations before analyses, then used a Gaussian distribution. For HR induction, we used a binomial distribution with the number of HR eggs considered as successes, and the total number of eggs as the total number of observations. For fruit damage, we used a beta distribution (Cribari-Neto & Zeileis, 2010).

For each phenotypic trait, we ran 4 different models with different genotype coding schemes and population structure controls. We ran two models coding genotypes differently: binary coding scheme: 0 = the plant had two reference alleles; 1 = the plant had at least one alternative allele, and dosage effect coding scheme: 0 = the plant had two reference alleles, 1 = the plant had one reference and one alternative allele, 2 = the plant had two alternative alleles. Coding genotypes as (0,1) can model dominance effects better, whereas coding genotypes as (0,1,2) can model dosage effects better. We also ran models with two different population structure controls: 1) treatment was used as a fixed effect, and replicate was used as a random effect, and 2) PC1 and PC2 from a genetic principal component analysis, which included all plants used in the GWAS runs, were included as covariates. As the only exception, treatment and replicate were listed as fixed effects when examining fruit damage. We did this because replicate had only two factor levels, which were too few to estimate random effects in the beta regression model (Cribari-Neto & Zeileis, 2010). Finally, within each model type, we used the *p.adjust* function in *R* with the FDR method to correct for multiple testing, and we considered associations with an FDR-adjusted *p*-value < 0.05 to be significant.

We report significant associations with two approaches: a more conservative approach and a less conservative approach. For the more conservative approach, we considered genotype-phenotype associations to be significant if associations for a given genomic region were significant in both models with different population structure controls (i.e., associations using both treatment/replicate and genetic PC1/PC2 were significant). For the less conservative approach, we considered genotype-phenotype associations to be significant if associations were significant in only one model (using either treatment/replicate or PC1/PC2). The rationale for reporting associations with two approaches is twofold: 1) using PCs to account for population structure may reduce power if the focal variant in genotype-phenotype associations load strongly onto the PCs (Privé et al., 2020; Zou et al., 2010) and 2), because there was no gene flow between plants from different treatment-replicate combinations and because all plants started from one starting population with the same pool of standing genetic variation, using the plants’ treatment and replicate can adequately control for population structure. However, using PCs is generally effective, and thus, to be more conservative, we report associations with two approaches.

#### 2.4.4 Candidate genes

To identify candidate genes associated with adaptation, we intersected genomic regions under selection with gene files from the reference genome used for alignment. We then assessed the gene function of associated *Arabidopsis thaliana* orthologs (https://phytozome-next.jgi.doe.gov/info/BrapaFPsc_v1_3).

#### 2.4.5 Genomic evolution of attraction and defense

To assess whether the strength of genomic evolution for attraction and defense traits differed between plants that evolved in different treatments, we first pruned traits with strong correlations (*r* > 0.7), leaving a total of 29 phenotypic traits. With this threshold, we removed the following volatile traits: total volatile emission, *(Z,Z)*-α-farnesene, and 2-aminobenzaldehyde. We also removed the following glucosinolate traits: total glucosinolate concentration, glucobrassicanapin, hydroxyglucobrassicin, and glucoraphasatin. Thus, for the “attraction” trait class (17 traits), we included morphological variables (plant height, the number of flowers and leaves, flower size), nectar, and floral volatile compounds (benzaldehyde, 1-butene-4-isothiocyanate, methyl benzoate, phenylethyl alcohol, *p*-anisaldehyde, methyl anthranilate, 3-hexenyl acetate, phenylacetaldehyde, benzyl nitrile, methyl salicylate, indole, *(E,E)*-α-farnesene). For the “defense” trait class (12 traits), we included the hypersensitive response (HR: the number of eggs that had a black necrotic ring around them) and glucosinolate compounds (sinigrin, glucobrassicin, gluconasturtiin, glucoalyssin, gluconapin, epiprogoitrin, progoitrin, glucoraphanin, glucoerucin, neoglucobrassicin, methoxyglucobrassicin). Similar to genotype-phenotype associations described above, we logged the emission values of all floral volatile traits and concentrations of glucosinolates before genome-wide association studies (GWAS) runs.

We then ran GWAS on the 29 phenotypic traits with EMMAX (Kang et al., 2010). For all GWAS runs, we included plants that coevolved with butterflies in both temperature environments and in the presence or absence of bumblebees. We also included plants from the control treatment and plants that evolved in response to non-coevolving butterflies (Figueira et al., *in review*). The average sample size per GWAS run ± standard deviation was 523.03 ± 66.09. We accounted for population structure by using kinship matrices generated with EMMAX (Kang et al., 2010). The number of eggs received by a plant was used as a covariate for the GWAS run on HR induction. After GWAS runs, we used the local score approach to identify genomic regions with significant phenotypic associations (Fariello et al., 2017).

We then calculated the change in allele frequencies (AF) for plants from each treatment as the AF of plants from the last generation minus the AF of plants from the first generation (G8-G1). To get the AF change associated with phenotypic traits, we only used SNPs within a genomic region with a significant phenotypic association and with a GWAS *p*-value < 0.0001. We took the AF change for these SNPs and subtracted them from the average AF change per chromosome. This was done to account for chromosomal differences in genetic evolution. We calculated the AF change per chromosome by using all SNPs at least 5kb away in either direction from any genomic regions with significant phenotypic association (results were qualitatively similar when using 10kb instead of 5kb). This was done to minimize correlations between average chromosomal AF changes and AF changes for genomic regions with phenotypic associations. After subtracting AF changes per SNP from the average chromosomal AF change, we calculated the average AF change per genomic region.

We then assessed whether AF changes underlying attraction and defense traits differed between plants that evolved in different treatments by running a generalized linear mixed model with the glmmTMB package (Brooks et al., 2017). The response variable was the absolute value of average AF changes for a given genomic region. We included temperature (ambient or elevated), bumblebee (presence or absence), and trait type (attraction or defense) as fixed effects. We also included the three-way interaction between temperature, bumblebee presence, and trait type, and all possible two-way interactions. We included replicate as a fixed effect because the model did not converge when replicate was included as a random effect. We included the genomic region ID (character string of: chromosome + bp of SNP with the lowest GWAS *p*-value) and phenotypic trait as random effects. Finally, values of the response variable fell within the range of [0,1), so we used an ordinal beta distribution (Kubinec, 2023).

## 3 Results

### 3.1 Genomic divergence

To examine whether the presence of bumblebees had a stronger effect on genomic divergence relative to elevated temperatures, we ran genome-wide *F_ST_* scans within replicates using 10kb windows. We quantified the effect of bumblebee presence on genomic divergence (bumblebee comparison) by running *F_ST_* scans between plants that coevolved with butterflies in the presence or absence of bumblebees (thus plants from different temperatures were grouped together), and we quantified the effect of elevated temperatures on genomic divergence (temperature comparison) by running *F_ST_* scans between plants that coevolved with butterflies in ambient or elevated temperature environments (thus plants from different bumblebee treatments were grouped together). On average, *F_ST_* values comparing the genomic divergence between plants coevolving in the presence or absence of bumblebees were more than double the *F_ST_* values comparing plants coevolving in different temperature environments (mean ± SEM; bumblebee presence: 0.0285 ± 0.0001; temperature: 0.0121 ± 0.0001; *df* = 1, *χ*^2^ = 12877, *p* > 0.001; Fig. 1A; Table S1). The bumblebee comparison had more 10kb windows with *F_ST_* values above 0.01, while the temperature comparison had more windows with *F_ST_* values within the range of 0 – 0.01 (Figs. 1A-1C).

**Figure 1.**
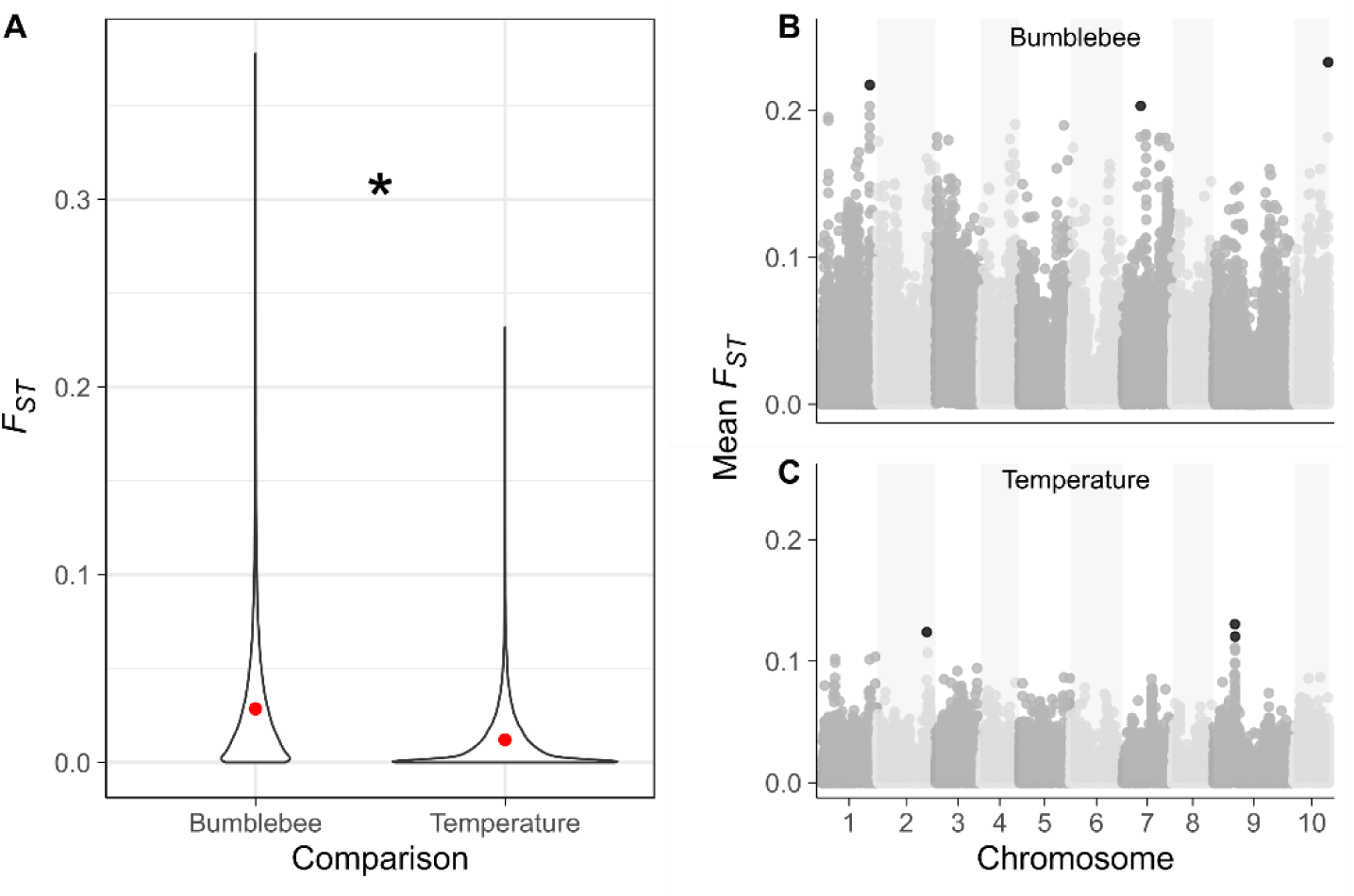
Genomic divergence is higher due to plants coevolving in the presence or absence of bumblebees rather than plants coevolving at different temperatures. (A) Violin plots show the distribution of *F_ST_* values from genome-wide *F_ST_* scans of 10kb windows. Wider edges of violin plots indicate a higher density of *F_ST_* values. Within violin plots for each comparison, the average *F_ST_* value is plotted as a red dot. (B, C) Manhattan plots of genome-wide *F_ST_* scans of 10kb windows are plotted for comparisons between (B) the presence/absence of bumblebees, and (C) ambient/elevated temperature environments. The y-axis shows the *F_ST_* value averaged across both replicates. Black points are windows with *F_ST_* values in the top 0.0001 percentile of *F_ST_* values within bumblebee or temperature comparisons. The x-axis shows the genomic regions of *B. rapa* with its 10 chromosomes.

We considered 10 kb windows as significant outliers when their average *F_ST_* value (mean across both replicates) fell in the top 0.0001 percentile (Figs. 1B, 1C); this threshold resulted in three outlier windows for both the bumblebee and temperature comparison (*F_ST_* range of outlier windows; bumblebee: 0.20 – 0.23; temperature: 0.12 – 0.13). We then identified candidate genes (*Arabidopsis thaliana* orthologs) that overlapped with outlier windows (Tables S2, S3). There were 5 candidate genes associated with the bumblebee comparison outlier windows, including potential ERF domain and F-box family proteins, and a gene associated with calmodulin binding (Table S2). There were also 5 candidate genes associated with the temperature comparison outlier windows and included a potential nodulin MtN21 /EamA-like transporter family protein, Trichome Birefringence-like 22, and phosphatidic acid phosphohydrolase 2 (Table S3).

### 3.2 Candidate genomic regions under selection

To understand how the presence/absence of bumblebees and temperature interacted to affect plant genomic adaptation in response to coevolution with butterflies, we first identified genomic regions exhibiting signatures of selection. We used Cochran–Mantel–Haenszel (CMH) tests, the local score approach, and the CLEAR method to find genomic regions with significant allele frequency changes between plants from the first (G1) and last generation (G8). We identified genomic regions with signatures of selection in plants from all treatments except for plants that coevolved with butterflies in elevated temperatures without bumblebees (Fig. 2; Tables S4-S6). Selected genomic regions were treatment-specific as they did not overlap between any of the treatments (Figs. 2B-E), and there were no overlaps between *F_ST_* outlier windows and treatment-specific selected genomic regions.

**Figure 2.**
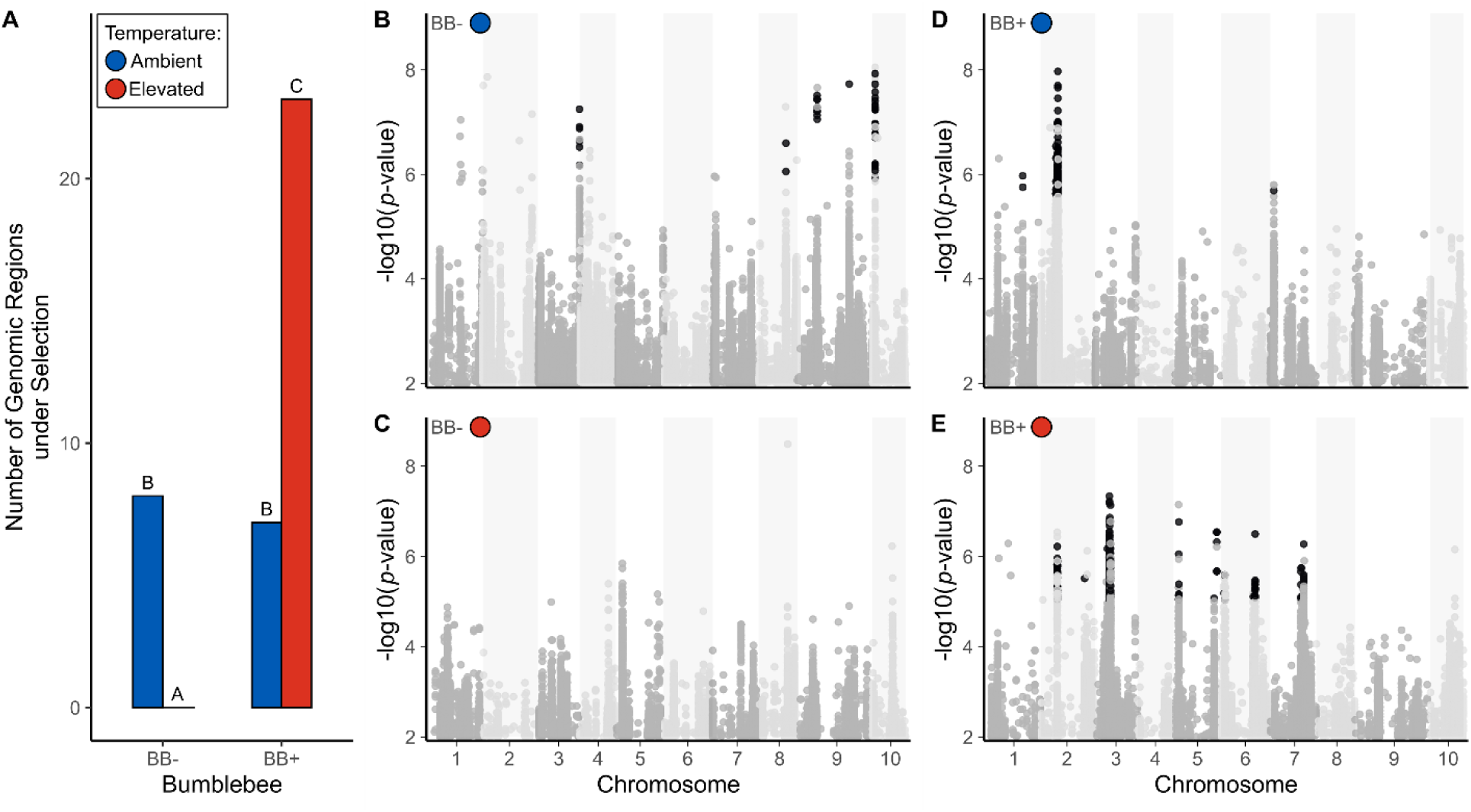
The presence of bumblebees and temperature interactively affect genomic adaptation for *Brassica rapa* plants coevolving with *Pieris rapae* butterflies. (A) The number of treatment-specific genomic regions with signatures of selection identified in genome-wide Cochran–Mantel–Haenszel (CMH) tests, the local score approach, and the CLEAR method are plotted as bars for plants that evolved in different treatments. Blue bars represent plants that coevolved in ambient temperatures (23±2°C), and red bars represent plants that coevolved in elevated temperatures (27±3°C with 24h of 30±3°C per week). The x-axis shows whether plants coevolved with butterflies in the absence of bumblebees (BB-) or in the presence of bumblebees (BB+). Differences in uppercase letters indicate significant pairwise differences between treatments in the proportion of treatment-specific selected genomic regions over the total number of selected genomic regions identified across all treatments. (B-E) Genome-wide scans for signatures of selection are represented using Manhattan plots. The -log10(*p*-value) from CMH tests is plotted along the y-axis. SNPs located within genomic regions exhibiting signatures of selection identified with CMH tests, the local score approach, and the CLEAR method are colored black. The x-axis shows the genomic coordinates of *B. rapa* with its 10 chromosomes. Manhattan plots are shown for plants from 4 treatments that coevolved with butterflies: (B) in ambient temperatures without bumblebees, (C) in elevated temperatures without bumblebees, (D) in ambient temperatures with bumblebees, and (E) in elevated temperatures with bumblebees.

Overall, we found an interactive effect of bumblebee presence and temperature on the number of selected genomic regions: the presence or absence of bumblebees affected the number of selected genomic regions differently depending on which temperature environment plants coevolved in (Fig 2A). More specifically, plants that coevolved in the absence of bumblebees had more genomic regions under selection when evolving in ambient relative to elevated temperatures (pairwise proportion test: *df* = 1, *χ*^2^ = 6.85, *p* = 0.013; Figs 2A-2C; Tables S4, S7). However, plants that coevolved in the presence of bumblebees had fewer genomic regions under selection when evolving in ambient temperatures rather than elevated temperatures (pairwise proportion test: *df* = 1, *χ*^2^ = 12.39, *p* = 0.001; Figs 2A, 2D-2E; Tables S5-S7). Comparing the number of selected genomic regions within temperature environments, plants that coevolved in ambient temperatures showed no difference in the number of selected genomic regions whether bumblebees were present or absent (pairwise proportion test: *df* = 1, *χ*^2^ = <0.01, *p* = 1; Figs 2A, 2B, 2D; Tables S4, S5, S7). Conversely, plants that coevolved in elevated temperatures had more selected genomic regions when bumblebees were present relative to when they were absent (pairwise proportion test: *df* = 1, *χ*^2^ = 12.39, *p* = 0.001; Figs 2A, 2C, 2E; Tables S6, S7). In summary, plant genomic adaptation in response to coevolution with butterflies was affected by different combinations of bumblebee presence and temperature.

To further examine the genomic basis of adaptation, we identified phenotypic and candidate gene associations within genomic regions that exhibited signatures of selection. We report genotype-phenotype associations using two approaches: 1) a more conservative approach - genotype-phenotype associations for a given genomic region were significant in two models with different population structure controls (associations using both treatment/replicate and genetic PC1/PC2 were significant), and 2) a less conservative approach - associations were significant in only one model (using either treatment/replicate or PC1/PC2).

For plants that coevolved with butterflies in ambient temperatures without bumblebees, we identified multiple genotype-phenotype associations and 7 candidate genes within selected genomic regions (Tables S4, S8-S11). With the more conservative approach for genotype-phenotype associations, we found associations with variation in the emission of phenylacetaldehyde and methyl salicylate, and an association with increased induction of the hypersensitive response (HR; Tables S8-S11). With the less conservative approach, we found associations with flower size and variation in several glucosinolate compounds such as glucoerucin, glucosbrassicin, neoglucobrassicin, hydroxyglucobrassicin, and glucobrassicanapin. Several of the 7 identified candidate genes had defense-related functions, such as a cysteine-rich RLK (receptor-like protein kinase) 8, a major latex-like protein 328, and a disease resistance-responsive (dirigent-like protein) family protein (Table S4). Additionally, we found a candidate potassium channel beta subunit 1 gene within the genomic region associated with increased HR induction, suggesting a potential link with HR defense.

For plants that coevolved with butterflies in ambient temperatures with bumblebees, we identified several genotype-phenotype associations and 15 candidate genes within selected genomic regions (Table S5, S12-S15). With the more conservative approach, we found one association with variation in the number of leaves (Table S12-S15). With the less conservative approach, we found associations with variation in flower size, nectar amount, and emission of methyl salicylate and 1-butene-4-isothiocyanate. We also found associations with two glucosinolate metabolites, glucosbrassicin and hydroxyglucobrassicin. Candidate genes were associated with various molecular functions such as protein binding/modification, ATP binding, protein kinase activity, and oxidoreductase activity (Table S5).

For plants that coevolved with butterflies in elevated temperatures with bumblebees, we identified multiple genotype-phenotype associations and 35 candidate genes within selected genomic regions (Table S6, S16-S19). With the more conservative approach, we found an association with a higher number of flowers (Tables S16-S19). With the less conservative approach, we found associations with variation in the emission of phenylacetaldehyde and indole. We also found associations with variation in HR induction and multiple glucosinolate compounds: progoitrin, glucobrassicin, hydroxyglucobrassicin, glucoraphasatin, glucobrassicanapin, epiprogoitrin, glucoerucin, and the total glucosinolate concentration. There were strong phenotypic correlations (*r* > 0.7; see methods) between several glucosinolate compounds, likely explaining why selected genomic regions were often associated with variation in multiple glucosinolates. Candidate gene functions were varied and included protein binding/modification, ATP binding, and protein kinase activity (Table S6).

### 3.3 Genomic evolution underlying attraction and defense traits

Finally, to compare the strength of genomic evolution underlying attraction and defense traits across plants that coevolved in different environmental conditions, we performed genome-wide association studies (GWAS) and computed allele frequency changes (between generations 1 and 8) for associated genomic regions. We considered morphological variables (plant height, the number of flowers and leaves, and flower size), nectar, and floral volatile compounds as attraction traits (n = 17). Conversely, we considered the hypersensitive response (HR; the number of eggs per plant that had a black necrotic ring around them) and glucosinolate compounds as defense traits (n = 12). We found that the strength of allele frequency changes differed between attraction and defense traits depending on whether plants coevolved with butterflies in the presence or absence of bumblebees (presence of bumblebees * trait type: *df* = 1, *χ*^2^ = 7.45, *p* = 0.006; Figs 3; Table S20). More specifically, allele frequency changes were greater for genomic regions associated with defense traits when plants coevolved in the presence of bumblebees rather than in their absence (*df* = Inf, *z-ratio* = 2.21, *p* = 0.027). Allele frequency changes underlying attraction traits did not differ when plants coevolved in the absence or presence of bumblebees (*df* = Inf, *z-ratio* = −1.61, *p* = 0.108). Finally, we found that the strength of allele frequency changes did not differ between attraction and defense traits depending on whether plants coevolved in ambient or elevated temperature environments (temperature * trait type: *df* = 1, *χ*^2^ = 0.063, *p* = 0.801).

**Figure 3.**
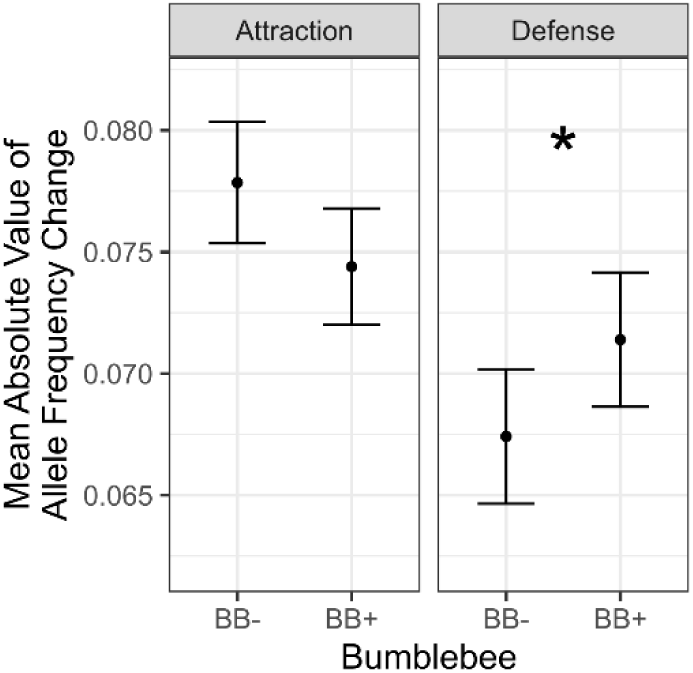
The strength of allele frequency changes differs between attraction and defense traits when plants coevolved with butterflies in the presence or absence of bumblebees. Average absolute values of allele frequency changes were calculated for genomic regions associated with phenotypic traits and plotted as black dots. 95% confidence intervals are plotted as error bars. The x-axis shows whether plants coevolved with butterflies in the absence of bumblebees (BB-) or in the presence of bumblebees (BB+). The * indicates significant differences between plants that evolved in different bumblebee treatments for defense traits.

## 4 Discussion

A major goal of the contemporary study of coevolution is to understand how genomic adaptation in response to coevolution is shaped by habitat context and global environmental change. To this end, we analyzed the genomes of plants from an experimental coevolution study (Rusman, Figueira, et al., 2025), where plants were allowed to coevolve with butterflies in the presence/absence of bumblebees and two temperature environments mirroring current climate and future climate predictions for Switzerland (Fischer et al., 2022). We show that additional pollination from non-coevolving bumblebees is a strong driver of genomic divergence and affects the genomic evolution underlying defense traits. In addition, we found strong non-additive interactive effects between bumblebee presence and temperature because the presence of non-coevolving bumblebees promoted genomic adaptation for coevolving plants, but only under elevated temperatures. Overall, our results show that variation in abiotic and biotic factors, including warmer temperatures mimicking global environmental change, can alter rapid adaptive genomic evolution as a consequence of coevolution.

### 4.1 Genomic divergence due to bumblebees and temperature

Variation in abiotic and biotic factors may affect the degree of genomic divergence between plant populations (Nosil, 2012; Nosil et al., 2009). In our study, we found that genomic divergence was larger between plants coevolving with butterflies in the presence or absence of bumblebees relative to plants coevolving with butterflies in different temperature environments (ambient/elevated). Both adaptation to different pollinators and temperature variation can promote genomic divergence (Figueira et al., 2025; Frachon et al., 2023; Wessinger et al., 2023; Xu et al., 2024), yet it was not shown before that effects from an additional pollinator can be larger relative to temperature-mediated effects. Two non-mutually exclusive mechanisms may have contributed to our finding of stronger pollinator-mediated genomic divergence. First, across all taxa (animals, bacteria, and plants), and in plants alone, biotic factors may have stronger fitness effects relative to abiotic factors (Briscoe Runquist et al., 2020), and adaptation in response to stronger selection imposed by bumblebees may have promoted more genomic divergence between our plant populations. Secondly, mutations with larger effect sizes may be more likely to be fixed in response to biotic rather than abiotic selection (Louthan & Kay, 2011). Because effect sizes can scale with the strength of selection, and stronger selection can promote genomic hitchhiking for neutral loci linked to selected loci (Nosil et al., 2009; Simons et al., 2018), large-effect mutations underlying biotic adaptation could drive greater genomic divergence. Regardless of the underlying mechanism, our results demonstrate that variation in the interacting community boosts genomic divergence between plant populations engaged in strict pairwise coevolution.

### 4.2 Genomic regions under selection

Variation in pollinator-mediated selection due to abiotic factors such as temperature may affect the strength of genomic adaptation (Figueira et al., 2025; Sletvold, 2019; Traine et al., 2024). Here, we found no genomic regions with signatures of selection for plants that coevolved with butterflies in elevated temperatures without bumblebees. Our results generally align with findings from two other studies: 1) a microbial study showing reduced coevolution at elevated temperatures (Duncan et al., 2017), and 2) a recent study examining plants grown from seeds from the same coevolution experiment as the current study (as well as non-coevolving plants not included here) which found no consistent genomic adaptation across replicates for plants that coevolved with butterflies in elevated temperatures (Ding & Schiestl, 2026). These genomic results may be explained by butterflies in elevated temperatures imposing reduced and/or more variable selection, potentially leading to the absence of consistent genomic adaptation. This hypothesis is supported by finding reduced reciprocal selection between plants and butterflies from the first generation of the current study in elevated relative to ambient temperatures (Rusman, Traine, et al., 2025). Additionally, when the same plants from the current study were phenotyped (Rusman, Figueira, et al., 2025), the authors found weaker selection on flower visitation (i.e., a less steep slope for the relationship between flower visitation and seed set) for plants that coevolved in elevated temperatures without bumblebees, relative to plants from other treatments. Overall, both mechanisms may have potentially contributed to relatively weak butterfly-mediated selection, ultimately resulting in the absence of consistent genomic adaptation based on SNP data.

Selection imposed by multiple factors, such as different pollinators, may act in the same direction to promote genomic adaptation (Burny et al., 2022; Wadgymar et al., 2022). Here, we found the most selected genomic regions for plants that coevolved with butterflies in elevated temperatures and the presence of bumblebees. One explanation that may partially account for the strong observed signal of genomic adaptation is that temperature-induced plasticity and synergistic selection from both butterflies and bumblebees promoted consistent genomic evolution. Support for this was found in the trait “number of flowers”, as earlier work on the same plant families used in the current study found that plants plastically produced more flowers when grown in elevated temperatures and that both butterflies and bumblebees imposed positive selection on the number of flowers (Rusman, Traine, et al., 2025; Traine et al., 2024, 2025). Furthermore, previous findings are consistent with phenotypic data measured on plants from the current study, as plants that coevolved with butterflies in elevated temperatures with bumblebees evolved the highest number of flowers relative to other coevolving plants and plants from the control treatment (Rusman, Figueira, et al., 2025). Finally, in the current study, we found a genotype-phenotype association for more flowers within a selected genomic region, thus providing support for selection patterns and phenotypic results at the genomic level.

### 4.3 Genomic evolution underlying defense traits

In the presence of additional pollinators, plants may become less dependent on mixed mutualistic-antagonistic pollinating herbivores for reproduction, making it adaptive to evolve increased defenses to reduce herbivory-mediated fitness costs (Kay & Anderson, 2025; Thompson & Cunningham, 2002). Here, we found that the genomic evolution underlying defense traits was higher for plants that coevolved with butterflies in the presence of bumblebees relative to plants that coevolved with butterflies in the absence of bumblebees. Our results align with phenotypic data measured on the same plants in the current study (Rusman, Figueira, et al., 2025), as plants coevolving with butterflies in ambient temperatures evolved increases in more defense traits when bumblebees were present relative to when bumblebees were absent. In addition, our results provide experimental support at the genomic level for findings from classic field studies examining the coevolutionary interaction between *Greya* moth species and *Lithophragma parviflorum* plants (Thompson & Cunningham, 2002; Thompson & Pellmyr, 1992). While the finding that bumblebees promoted plant genomic evolution underlying defense traits is statistically significant, it is important to acknowledge that the effect size is quite small (∼1% difference in allele frequency changes). Still, our results broadly show that novel interactions with mutualistic pollinators shape the genomic evolution underlying traits associated with defensive functions, in addition to well-known effects on mutualistic attraction traits (Bradshaw et al., 1998).

### 4.4 Conclusions and outlook

Coevolutionary interactions are pervasive across the tree of life, and the genomic consequences of such interactions are affected by abiotic and biotic factors (Thompson, 2005). In the current study, we found that bumblebees seemingly promoted stronger genomic adaptation in elevated temperatures, suggesting that novel biotic interactions due to range shifts resulting from global warming may facilitate adaptation to potential coevolutionary interactions. Overall, we show that the interacting community outside of strict pairwise coevolution shapes genomic trajectories and will be of key importance in determining how plants adapt to novel conditions in the face of ongoing global environmental change.

## Author Contributions

Conceptualization: F.P.S., T.F., Q.R. Experimental work: T.F., Q.R. Analysis: T.F. Writing: T.F., F.P.S., Q.R.

## Acknowledgments

We thank Corinne Hertäg for help during the floral scent analyses and quantification, Laura Dällenbach, Basil Züllig, Chiara Joy Caspers, and Lena Schneider for help with various experimental tasks, Rayko Jonas and Markus Meierhofer for taking care of the plants, Juan Traine for help in creating and maintaining experimental plant and butterfly lines, Giacomo Potente and Yanqian Ding for help with genome assembly pipelines, and the Genetic Diversity Center of Zurich for help with DNA extractions. The research was funded by the Swiss National Science Funds (SNF grant no. 310030_208158) to F.P.S.

## Data Accessibility Statement

The raw data used in this publication is published in a previous study. The raw phenotypic data is available on figshare (doi: 10.6084/m9.figshare.31134943). Raw genetic reads were published on ENA with accession number: PRJEB106979. The data to produce figures and code will be made available on figshare (doi: 10.6084/m9.figshare.31134943) upon acceptance of the manuscript and can be accessed by reviewers(https://figshare.com/s/425e13fc446f4f9db743).

