## Supplemental Information for "Abiotic and biotic ecological factors affect the genomic consequences of plant-butterfly coevolution"

Tyler Figueira, Quint Rusman, Florian P. Schiestl*
Department of Systematic and Evolutionary Botany, University of Zürich
*Corresponding author

Table S1. Bumblebee presence influences genetic differentiation more than temperature. $F_{ST}$ scans of 10 kb windows were performed between experimental plants from the final generation (G8), and then a generalized linear mixed model was run with $F_{ST}$ values as the response variable ($N$ = 110667). The only fixed effect was the comparison term, which had two levels: 1) bumblebee: presence or absence, and 2) temperature: ambient or elevated. Replicate, chromosome, and start position of 10 kb window nested within the chromosome were included as random factors. An ordinal beta distribution was used as $F_{ST}$ values fell within the range of [0,1).

| Term | $\chi^2$ | *df* | *p*-value |
|---|---|---|---|
| Comparison | 16902 | 1 | **< 0.001** |

Table S2. Candidate genes associated with the bumblebee comparison $F_{ST}$ outlier windows. We considered 10 kb windows as significant outliers when their average $F_{ST}$ (across both replicates) fell in the top 0.0001 percentile. The column descriptions are as follows: CHROM = chromosome, BIN_START = starting base position of the $F_{ST}$ window, BIN_END = end base position of the $F_{ST}$ window, WEIGHTED_FST = the average $F_{ST}$ across both replicates for the window. The following columns describe potential functions and associations with the candidate genes: Pfam, Panther, KOG, KEGG/ec, KO, GO. Best-hit-arabi-name and arabi-defline indicate the best hit gene name and annotation based on *Arabidopsis thaliana*. The table has too many columns to fit onto this page and is given as a separate file.

Table S3. Candidate genes associated with temperature comparison $F_{ST}$ outlier windows. We considered 10 kb windows as significant outliers when their average $F_{ST}$ (across both replicates) fell in the top 0.0001 percentile. The column descriptions are as follows: CHROM = chromosome, BIN_START = starting base position of the $F_{ST}$ window, BIN_END = end base position of the $F_{ST}$ window, WEIGHTED_FST = the average $F_{ST}$ across both replicates for the window. The following columns describe potential functions and associations with the candidate genes: Pfam, Panther, KOG, KEGG/ec, KO, GO. Best-hit-arabi-name and arabi-defline indicate the best hit gene name and annotation based on *Arabidopsis thaliana*. The table has too many columns to fit onto this page and is given as a separate file.

Table S4. Genomic regions that exhibit signatures of selection are shown for plants that coevolved with butterflies in ambient temperatures in the absence of bumblebees. The start and end region columns show the boundaries of the selected regions. The most significant SNP column indicates the SNP with the lowest *p*-value in Cochran–Mantel–Haenszel (CMH) tests within the selected region. The candidate genes column shows which *Arabidopsis Thaliana* orthologs overlap with selected genomic regions.

| Chromosome | Start Region | End Region | Most Significant SNP | Candidate Genes |
|---|---|---|---|---|
| 3 | 24912162 | 24914005 | 3_24912963 | Cysteine-rich RLK (RECEPTOR-like protein kinase) 8 |
| 3 | 24914973 | 24918177 | 3_24917717 | |
| 8 | 16324792 | 16345782 | 8_16340487 | MLP-like protein 328, Disease resistance-responsive (dirigent-like protein) family protein, cation/H+ exchanger 17 |
| 9 | 11758282 | 11762414 | 9_11761431 | Esterase/lipase/thioesterase family protein |
| 9 | 30966399 | 30970392 | 9_30970098 | |
| 10 | 1531291 | 1532593 | 10_1531886 | Lipoyltransferase 2 |
| 10 | 1536682 | 1538028 | 10_1537192 | |
| 10 | 1572589 | 1574028 | 10_1573760 | Potassium channel beta subunit 1 |

Table S5. Genomic regions that exhibit signatures of selection are shown for plants that coevolved with butterflies in ambient temperatures in the presence of bumblebees. The start and end region columns show the boundaries of the selected regions. The most significant SNP column indicates the SNP with the lowest *p*-value in CMH tests within the selected region. The candidate genes column shows which *Arabidopsis Thaliana* orthologs overlap with selected genomic regions.

| Chromosome | Start Region | End Region | Most Significant SNP | Candidate Genes |
|---|---|---|---|---|
| 1 | 20254233 | 20343159 | 1_20255677 | F-box/RNI-like superfamily protein, NSP-interacting kinase 3, Ulp1 protease family protein, D-alanine--D-alanine ligase family, U2 snrnp auxiliary factor, large subunit, splicing factor, sulfotransferase 17, SNF1-related protein kinase 2.10 |
| 2 | 9080907 | 9084242 | 2_9081986 | |

| | | | | |
|---|---|---|---|---|
| 2 | 9089564 | 9091183 | 2_9090806 | Ubiquitin carboxyl-terminal hydrolase-related protein, Ubiquitin carboxyl-terminal hydrolase family protein |
| 2 | 9810109 | 9860808 | 2_9811118 | Ribulose bisphosphate carboxylase (small chain) family protein, LOB domain-containing protein 40, cytochrome P450, family 735, subfamily A, polypeptide 2 |
| 2 | 9895248 | 9906533 | 2_9905648 | |
| 2 | 10027221 | 10028802 | 2_10027384 | |
| 2 | 10101372 | 10104223 | 2_10104223 | ARM repeat superfamily protein |
| 2 | 10255500 | 10268308 | 2_10261818 | Protein of unknown function (DUF707), glycine-rich protein |
| 7 | 2087830 | 2095569 | 7_2088831 | SET domain group 37 |

Table S6. Genomic regions that exhibit signatures of selection are shown for plants that coevolved with butterflies in elevated temperatures in the presence of bumblebees. The start and end region columns show the boundaries of the selected regions. The most significant SNP column indicates the SNP with the lowest *p*-value in CMH tests within the selected region. The candidate genes column shows which *Arabidopsis Thaliana* orthologs overlap with selected genomic regions.

| Chromosome | Start Region | End Region | Most Significant SNP | Candidate Genes |
|---|---|---|---|---|
| 2 | 9797581 | 9800150 | 2_9798581 | Mannose-6-phosphate isomerase, type I |
| 2 | 9911712 | 9932429 | 2_9912712 | Zinc finger (C3HC4-type RING finger) family protein / BRCT domain-containing protein, F-box and associated interaction domains-containing protein |
| 2 | 26320223 | 26321745 | 2_26320683 | |
| 3 | 7163822 | 7167050 | 3_7166785 | Pseudouridine synthase family protein, S-adenosyl-L-methionine-dependent methyltransferases superfamily protein |
| 3 | 8328894 | 8334190 | 3_8333838 | Protein of unknown function (DUF3527), Caleosin-related family protein |
| 3 | 8350472 | 8358079 | 3_8351812 | RNA-binding (RRM/RBD/RNP motifs) family protein |

| | | | | |
|---|---|---|---|---|
| 3 | 8405034 | 8407647 | 3_8407647 | P-loop containing nucleoside triphosphate hydrolases superfamily protein, Galactose oxidase/kelch repeat superfamily protein |
| 3 | 8419370 | 8453443 | 3_8420413 | Zinc finger C-x8-C-x5-C-x3-H type family protein, HAD superfamily, subfamily IIIB acid phosphatase, RAB gtpase homolog A1H, homeobox-3, Calcium-binding EF-hand family protein |
| 3 | 8578407 | 8590234 | 3_8588198 | Embryo defective 2739, myosin heavy chain-related, Protein phosphatase 2C family protein, RNA polymerase I specific transcription initiation factor RRN3 protein |
| 3 | 8686724 | 8688398 | 3_8688188 | EYES ABSENT homolog |
| 3 | 8958838 | 8965426 | 3_8959838 | Receptor-like kinase in flowers 3, pleiotropic drug resistance 6 |
| 3 | 8965377 | 8967771 | 3_8965426 | |
| 3 | 8975023 | 8989493 | 3_8977596 | Family of unknown function (DUF662) |
| 3 | 9013154 | 9020481 | 3_9020470 | Plant protein of unknown function (DUF868) |
| 3 | 9021032 | 9023093 | 3_9022032 | |
| 3 | 9042409 | 9043986 | 3_9043409 | Ribosomal protein L12-B, phloem protein 2-A15 |
| 5 | 3035558 | 3041711 | 5_3040677 | Eukaryotic translation initiation factor 2 subunit 1 |
| 5 | 3148947 | 3159386 | 5_3150649 | AP2/B3-like transcriptional factor family protein, Glutathione S-transferase, C-terminal-like; Translation elongation factor EF1B/ribosomal protein S6, Phosphoinositide-specific phospholipase C family protein, Pollen Ole e 1 allergen and extensin family protein |
| 5 | 24250346 | 24253146 | 5_24251433 | |
| 5 | 25836666 | 25837886 | 5_25837727 | Leucine-rich repeat protein kinase family protein |
| 6 | 1816538 | 1817574 | 6_1817538 | Prolyl oligopeptidase family protein |
| 6 | 2216641 | 2218044 | 6_2217855 | Stomatal cytokinesis defective / SCD1 protein (SCD1) |
| 6 | 19249322 | 19262372 | 6_19260337 | NAC domain-containing protein 102, RNA-directed DNA polymerase (reverse |

| | | | | |
|---|---|---|---|---|
| | | | | transcriptase)-related family protein, Glycosyl hydrolase family 35 protein |
| 6 | 20241964 | 20250275 | 6_20247112 | |
| 6 | 20290801 | 20295146 | 6_20294320 | Protein kinase superfamily protein, |
| 6 | 20312924 | 20314691 | 6_20314691 | Concanavalin A-like lectin protein kinase family protein |
| 6 | 20314831 | 20317192 | 6_20316348 | |
| 7 | 18295123 | 18296254 | 7_18295942 | H(+)-atpase 9 |
| 7 | 18297435 | 18298483 | 7_18298435 | Transducin/WD40 repeat-like superfamily protein |
| 7 | 18298763 | 18301022 | 7_18299021 | Suppressor of auxin resistance 3 |
| 7 | 18444346 | 18445730 | 7_18445612 | Protein kinase superfamily protein with octicosapeptide/Phox/Bem1p domain |
| 7 | 18445975 | 18450978 | 7_18450235 | FTSH protease 12 |
| 7 | 19902967 | 19907221 | 7_19904674 | SNF7 family protein |
| 7 | 19970016 | 19976037 | 7_19971115 | Hydroxyproline-rich glycoprotein family protein |
| 7 | 19993218 | 19994565 | 7_19993779 | ELF4-like 2, alpha/beta-Hydrolases superfamily protein |

Table S7. Pairwise differences between treatments illustrate how temperature and bumblebee presence affect the number of genomic regions under selection. Pairwise tests were run between plants from all treatments (except for control treatments) comparing the proportion of: treatment-specific genomic regions / total number of genomic regions identified across all treatments. We corrected for multiple testing with the false discovery rate (FDR). Plant evolutionary treatments are as follows: (Ambient BB-) = plants coevolved in ambient temperatures without bumblebees; (Elevated BB-) = plants coevolved in elevated temperatures without bumblebees; (Ambient BB+) = plants coevolved in ambient temperatures with bumblebees; (Elevated BB+) = plants coevolved in elevated temperatures with bumblebees.

| Comparison | $\chi^2$ | *df* | *p*-value | FDR(*p*-value) |
|---|---|---|---|---|
| (Ambient BB+) – (Ambient BB-) | < 0.01 | 1 | 1 | 1 |
| (Ambient BB+) – (Elevated BB+) | 12.39 | 1 | **< 0.001** | **0.001** |
| (Ambient BB+) – (Elevated BB-) | 5.66 | 1 | **0.017** | **0.021** |
| (Ambient BB-) – (Elevated BB+) | 10.68 | 1 | **0.001** | **0.002** |
| (Ambient BB-) – | 6.85 | 1 | **0.009** | **0.013** |

| | | | | |
|---|---|---|---|---|
| (Elevated BB-) | | | | |
| (Elevated BB+) – (Elevated BB-) | 30.18 | 1 | **< 0.001** | **< 0.001** |

Tables S8-S11: Genotype-phenotype associations for plants that coevolved with butterflies in ambient temperatures without bumblebees.

Table S8. For plants that coevolved with butterflies in ambient temperatures without bumblebees, phenotypic associations with genomic regions under selection using genotypes coded as (0,1) and treatment and replicate to control for population structure are listed. Genotypes were coded as (0,1), plant genotypes were coded binarily depending on whether they differed from the reference allele (both alleles were identical to the reference = 0; any one or both of the two alleles were different relative to the reference = 1). We ran the following generalized linear mixed model: Trait ~ plant genotype (0,1) + (1|replicate) + treatment.

The columns are as follows: name_2 lists which phenotypic trait was tested, the estimate gives the estimated effect size, chisq gives the $\chi^2$ using the ANOVA function from the car package in R, n gives the sample size for the model, the SNP column shows the SNP that genotypes were used to test for associations and fdr_p gives the *p*-value corrected with the FDR method.  We considered phenotypic traits with a fdr_p > 0.05 to be significant associations. The table is larger than 100 rows and is given as a separate file.

Table S9. For plants that coevolved with butterflies in ambient temperatures without bumblebees, phenotypic associations with genomic regions under selection using genotypes coded as (0,1) and PC1 and PC2 to control for population structure are listed. Genotypes were coded as (0,1), plant genotypes were coded binarily depending on whether they differed from the reference allele (both alleles were identical to the reference = 0; any one or both of the two alleles were different relative to the reference = 1). We ran the following generalized linear mixed model: Trait ~ plant genotype (0,1) + PC1 + PC2.

The columns are as follows: name_2 lists which phenotypic trait was tested, the estimate gives the estimated effect size, chisq gives the $\chi^2$ using the ANOVA function from the car package in R, n gives the sample size for the model, the SNP column shows the SNP that genotypes were used to test for associations and fdr_p gives the *p*-value corrected with the FDR method.  We considered phenotypic traits with a fdr_p > 0.05 to be significant associations. The table is larger than 100 rows and is given as a separate file.

Table S10. For plants that coevolved with butterflies in ambient temperatures without bumblebees, phenotypic associations with genomic regions under selection using genotypes coded as (0,1,2) and treatment and replicate to control for population structure are listed. Genotypes were coded as (0,1,2): a genotype value of 0 indicates both alleles were the reference

allele, a genotype value of 1 indicates one allele was the reference and the other was the alternative, and a genotype value of 2 indicates that both alleles were the alternative allele.
We ran the following generalized linear mixed model: Trait ~ plant genotype (0,1,2) + (1|replicate) + treatment.

The columns are as follows: name_2 lists which phenotypic trait was tested, the estimate gives the estimated effect size, chisq gives the $\chi^2$ using the ANOVA function from the car package in R, n gives the sample size for the model, the SNP column shows the SNP that genotypes were used to test for associations and fdr_p gives the *p*-value corrected with the FDR method. We considered phenotypic traits with a fdr_p > 0.05 to be significant associations. The table is larger than 100 rows and is given as a separate file.

Table S11. For plants that coevolved with butterflies in ambient temperatures without bumblebees, phenotypic associations with genomic regions under selection using genotypes coded as (0,1,2) and PC1 and PC2 to control for population structure are listed. Genotypes were coded as (0,1,2): a genotype value of 0 indicates both alleles were the reference allele, a genotype value of 1 indicates one allele was the reference and the other was the alternative, and a genotype value of 2 indicates that both alleles were the alternative allele.
We ran the following generalized linear mixed model: Trait ~ plant genotype (0,1,2) + PC1 + PC2.

The columns are as follows: name_2 lists which phenotypic trait was tested, the estimate gives the estimated effect size, chisq gives the $\chi^2$ using the ANOVA function from the car package in R, n gives the sample size for the model, the SNP column shows the SNP that genotypes were used to test for associations and fdr_p gives the *p*-value corrected with the FDR method. We considered phenotypic traits with a fdr_p > 0.05 to be significant associations. The table is larger than 100 rows and is given as a separate file.

Tables S12-S15: Genotype-phenotype associations for: plants that coevolved with butterflies in ambient temperatures with bumblebees

Table S12. For plants that coevolved with butterflies in ambient temperatures with bumblebees, phenotypic associations with genomic regions under selection using genotypes coded as (0,1) and treatment and replicate to control for population structure are listed. Genotypes were coded as (0,1), plant genotypes were coded binarily depending on whether they differed from the reference allele (both alleles were identical to the reference = 0; any one or both of the two alleles were different relative to the reference = 1). We ran the following generalized linear mixed model: Trait ~ plant genotype (0,1) + (1|replicate) + treatment.

The columns are as follows: name_2 lists which phenotypic trait was tested, the estimate gives the estimated effect size, chisq gives the $\chi^2$ using the ANOVA function from the car package in R, n gives the sample size for the model, the SNP column shows the SNP that genotypes were used to test for associations and fdr_p gives the *p*-value corrected with the FDR method. We considered phenotypic traits with a fdr_p > 0.05 to be significant associations. The table is larger than 100 rows and is given as a separate file.

Table S13. For plants that coevolved with butterflies in ambient temperatures with bumblebees, phenotypic associations with genomic regions under selection using genotypes coded as (0,1) and PC1 and PC2 to control for population structure are listed. Genotypes were coded as (0,1), plant genotypes were coded binarily depending on whether they differed from the reference allele (both alleles were identical to the reference = 0; any one or both of the two alleles were different relative to the reference = 1). We ran the following generalized linear mixed model: Trait ~ plant genotype (0,1) + PC1 + PC2.

The columns are as follows: name_2 lists which phenotypic trait was tested, the estimate gives the estimated effect size, chisq gives the $\chi^2$ using the ANOVA function from the car package in R, n gives the sample size for the model, the SNP column shows the SNP that genotypes were used to test for associations and fdr_p gives the *p*-value corrected with the FDR method. We considered phenotypic traits with a fdr_p > 0.05 to be significant associations. The table is larger than 100 rows and is given as a separate file.

Table S14. For plants that coevolved with butterflies in ambient temperatures with bumblebees, phenotypic associations with genomic regions under selection using genotypes coded as (0,1,2) and treatment and replicate to control for population structure are listed. Genotypes were coded as (0,1,2): a genotype value of 0 indicates both alleles were the reference allele, a genotype value of 1 indicates one allele was the reference and the other was the alternative, and a genotype value of 2 indicates that both alleles were the alternative allele.
We ran the following generalized linear mixed model: Trait ~ plant genotype (0,1,2) + (1|replicate) + treatment.

The columns are as follows: name_2 lists which phenotypic trait was tested, the estimate gives the estimated effect size, chisq gives the $\chi^2$ using the ANOVA function from the car package in R, n gives the sample size for the model, the SNP column shows the SNP that genotypes were used to test for associations and fdr_p gives the *p*-value corrected with the FDR method. We considered phenotypic traits with a fdr_p > 0.05 to be significant associations. The table is larger than 100 rows and is given as a separate file.

Table S15. For plants that coevolved with butterflies in ambient temperatures with bumblebees, phenotypic associations with genomic regions under selection using genotypes coded as (0,1,2) and PC1 and PC2 to control for population structure are listed. Genotypes were coded as (0,1,2): a genotype value of 0 indicates both alleles were the reference allele, a genotype value of 1 indicates one allele was the reference and the other was the alternative, and a genotype value of 2 indicates that both alleles were the alternative allele.
We ran the following generalized linear mixed model: Trait ~ plant genotype (0,1,2) + PC1 + PC2.

The columns are as follows: name_2 lists which phenotypic trait was tested, the estimate gives the estimated effect size, chisq gives the $\chi^2$ using the ANOVA function from the car package in R, n gives the sample size for the model, the SNP column shows the SNP that genotypes were used to test for associations and fdr_p gives the *p*-value corrected with the FDR method. We considered phenotypic traits with a fdr_p > 0.05 to be significant associations. The table is larger than 100 rows and is given as a separate file.

Tables S16-S19: Genotype-phenotype associations for: plants that coevolved with butterflies in elevated temperatures with bumblebees

Table S16. For plants that coevolved with butterflies in elevated temperatures with bumblebees, phenotypic associations with genomic regions under selection using genotypes coded as (0,1) and treatment and replicate to control for population structure are listed. Genotypes were coded as (0,1), plant genotypes were coded binarily depending on whether they differed from the reference allele (both alleles were identical to the reference = 0; any one or both of the two alleles were different relative to the reference = 1). We ran the following generalized linear mixed model: Trait ~ plant genotype (0,1) + (1|replicate) + treatment.

The columns are as follows: name_2 lists which phenotypic trait was tested, the estimate gives the estimated effect size, chisq gives the $\chi^2$ using the ANOVA function from the car package in R, n gives the sample size for the model, the SNP column shows the SNP that genotypes were used to test for associations and fdr_p gives the *p*-value corrected with the FDR method. We considered phenotypic traits with a fdr_p > 0.05 to be significant associations. The table is larger than 100 rows and is given as a separate file.

Table S17. For plants that coevolved with butterflies in elevated temperatures with bumblebees, phenotypic associations with genomic regions under selection using genotypes coded as (0,1) and PC1 and PC2 to control for population structure are listed. Genotypes were coded as (0,1), plant genotypes were coded binarily depending on whether they differed from the reference allele (both alleles were identical to the reference = 0; any one or both of the two alleles were different relative to the reference = 1). We ran the following generalized linear mixed model: Trait ~ plant genotype (0,1) + PC1 + PC2.

The columns are as follows: name_2 lists which phenotypic trait was tested, the estimate gives the estimated effect size, chisq gives the $\chi^2$ using the ANOVA function from the car package in R, n gives the sample size for the model, the SNP column shows the SNP that genotypes were used to test for associations and fdr_p gives the *p*-value corrected with the FDR method. We considered phenotypic traits with a fdr_p > 0.05 to be significant associations. The table is larger than 100 rows and is given as a separate file.

Table S18. For plants that coevolved with butterflies in elevated temperatures with bumblebees, phenotypic associations with genomic regions under selection using genotypes coded as (0,1,2)

and treatment and replicate to control for population structure are listed. Genotypes were coded as (0,1,2): a genotype value of 0 indicates both alleles were the reference allele, a genotype value of 1 indicates one allele was the reference and the other was the alternative, and a genotype value of 2 indicates that both alleles were the alternative allele.
We ran the following generalized linear mixed model: Trait ~ plant genotype (0,1,2) + (1|replicate) + treatment.

The columns are as follows: name_2 lists which phenotypic trait was tested, the estimate gives the estimated effect size, chisq gives the $\chi^2$ using the ANOVA function from the car package in R, n gives the sample size for the model, the SNP column shows the SNP that genotypes were used to test for associations and fdr_p gives the *p*-value corrected with the FDR method. We considered phenotypic traits with a fdr_p > 0.05 to be significant associations. The table is larger than 100 rows and is given as a separate file.

Table S19. For plants that coevolved with butterflies in elevated temperatures with bumblebees, phenotypic associations with genomic regions under selection using genotypes coded as (0,1,2) and PC1 and PC2 to control for population structure are listed. Genotypes were coded as (0,1,2): a genotype value of 0 indicates both alleles were the reference allele, a genotype value of 1 indicates one allele was the reference and the other was the alternative, and a genotype value of 2 indicates that both alleles were the alternative allele.
We ran the following generalized linear mixed model: Trait ~ plant genotype (0,1,2) + PC1 + PC2.

The columns are as follows: name_2 lists which phenotypic trait was tested, the estimate gives the estimated effect size, chisq gives the $\chi^2$ using the ANOVA function from the car package in R, n gives the sample size for the model, the SNP column shows the SNP that genotypes were used to test for associations and fdr_p gives the *p*-value corrected with the FDR method. We considered phenotypic traits with a fdr_p > 0.05 to be significant associations. The table is larger than 100 rows and is given as a separate file.

Table S20. The magnitude of allele frequency changes differs between attraction and defense traits when plants evolve in the presence or absence of bumblebees. A generalized linear mixed model was run with the absolute value of allele frequency changes per genomic region as the response variable ($N$ = 12536). Fixed effects included: temperature (ambient or elevated), bumblebee (presence or absence), and trait type (attraction or defense). We also included the three-way interaction between temperature, bumblebee presence, and trait type, and all possible two-way interactions. Replicate was included as a fixed effect as well. The region ID (character string of: chromosome + bp of SNP with the lowest GWAS *p*-value) and trait were included as random effects. An ordinal beta distribution was used as the response variable fell within the range of [0,1).

| Term | $\chi^2$ | *df* | *p*-value |
|---|---|---|---|

| | | | |
|---|---|---|---|
| Temperature | 0.49 | 1 | 0.486 |
| Bumblebee presence | 0.06 | 1 | 0.8 |
| Trait type | 0.18 | 1 | 0.667 |
| Replicate | 0.13 | 1 | 0.728 |
| Temperature* Bumblebee presence | 0.74 | 1 | 0.391 |
| Temperature* Trait type | 0.06 | 1 | 0.801 |
| Bumblebee presence * Trait type | 7.45 | 1 | **0.006** |
| Temperature* Bumblebee presence* Trait type | 0.35 | 1 | 0.555 |
